# FAIR Data Station for Lightweight Metadata Management & Validation of Omics Studies

**DOI:** 10.1101/2022.08.03.502622

**Authors:** Bart Nijsse, Peter J. Schaap, Jasper J. Koehorst

## Abstract

**Background:** The Life sciences is an interdisciplinary field of research and one of the the biggest suppliers of scientific data. Reusing and connecting this data can uncover hidden insights and lead to new concepts, provided there is machine-actionable metadata available about the scientific experiments conducted with a degree of completeness that reflect the FAIR guiding principles. While stakeholders have embraced the FAIR principles, in practice there are a limited number of easy to adopt practical implementations available that fulfil the needs of data producers.

**Findings:** We developed the FAIR Data Station, a lightweight application written in Java, that aims to support researchers in managing research metadata according to the FAIR principles. It uses the ISA metadata framework and metadata standards to capture experimental metadata. The FAIR Data Station metadata registration workflow consists of three main modules. Based on the minimal information checklist(s) selected by the user, a web-based “form generation module” creates a standardized metadata template Excel workbook which is used as a familiar environment for offline sample metadata registration. A web-based “validation module” checks the format of the metadata recorded in the workbook. The “resource module” subsequently exports the validated set of recorded metadata into an RDF data file, enabling (cross-project) meta data searches.

**Conclusions:** Turning FAIR into reality requires the availability of easy to adopt data FAIRification workflows that provide immediate beneficial incentives to the individual researcher. As such the FAIR Data Station provides in addition to the means to correctly FAIRify sequence data, the means to build searchable databases of (local) projects and can assists in ENA metadata submission of sequence data. The FAIR Data Station is available at http://fairbydesign.nl.

## Background

Online repositories sharing scientific data are vital for the advancement of science. Data sharing improves research transparency and promotes the validation of experimental methods and scientific conclusions. Data sharing enables data reuse and facilitates knowledge discovery using new analysis tools. Essential for reusing shared scientific data is the availability of machine-actionable metadata of sufficient quality about the scientific experiments conducted with a degree of completeness that reflects the FAIR guiding principles: Findable, Accessible, Interoperable, Reusable [1].

Several concepts have been developed to assist in the data FAIR-ification process. The ISA metadata framework standard [2] specifies an abstract model to capture experimental metadata using three core levels, Investigation, Study and Assay. The GO-FAIR initiative [3] suggests a seven-step workflow for data FAIRification. They, however, do not provide practical implementations of the technological components needed in the FAIRification process. This is because FAIR is not a standard, but a set of guidelines open to interpretation.

A key feature of properly FAIRified data is a high level of data Interoperability. From a data producer/user point of view two levels are important: structural and semantic interoperability. Structural interoperability defines the format of the data allowing the data to be interpreted by multiple systems or devices. The FASTA sequence format, for instance, is the most implemented data standard for sequence data [4, 5]. Semantic interoperability entails the transformation of ambiguous human-understandable metadata in a standardized machine-actionable open format. Creating semantic interoperability is complex and can require significant efforts. To ensure that sets of metadata is sufficient for the data to be unambiguously described, standardized minimal information models and checklists, detailing those reporting requirements, have been developed for wide array of experimental data [6].

The wide-spread adoption of high-throughput sequencing methods has made sequence data the major big data generator of the Life Sciences [7]. Sequence data is a special case as it implies a large-scale assessment of a single type of molecules. This property and its representation in standard FASTA format make the sequence data type an excellent candidate for data reuse. To assist in the FAIRification process of sequence data, the Genomic Standards Consortium [8] has developed a widely accepted family of minimum information standard checklists about any (x) Sequence (MIxS). While they were developed with sequence data in mind, these guidelines can also be used to describe sample metadata of other (Omics) studies

To help researchers to FAIRify their experimental data in line with accepted standards we have developed the FAIR Data Station (FAIR-DS). The overall goal of this lightweight stand-alone tool is to assist the domain researcher in creating high-quality FAIR metadata. The FAIR-DS supports metadata standards such as the MIxS set of standards implemented by the main sequence databases such as the European Nucleotide Archive (ENA), Genbank, MGnify (EBI Metagenomics), JGI-GOLD and others (see https://doi.org/10.25504/FAIRsharing.9aa0zp for more) and can be used to streamline metadata submission of sequence data to the ENA repository. The output of the FAIR Data Station can also be used to build a local metadata database of (similar) projects, while the default set of mandatory metadata fields can easily be expanded to align with the internal standards of a research group.

## Design considerations

For metadata registration the FAIR-DS uses an extended version of the original three-tier Investigation, Study, Assay (ISA) metadata framework [https://isa-tools.org]. The Investigation layer contains human readable project related metadata: title, authors and a minimal amount of high-level information for humans to understand the overall goals of the experiment. The Study layer describes a specific research line. As one Investigation can have several research lines, each Study layer has a unique identifier, a study title, and a description of the experimental design of the specific line of research. If the investigation involves only a single study, this title can be copied from the investigation layer. As an extension to the original three-tier ISA-model in between Study and Assay two additional layers of information were added: Observation unit and Sample. An Observation unit is an object that is subject to instances of observation and measurement [9] or the “thing” on which measurements are taken. The number of Observation units should be in line with the experimental design. The Sample layer describes the conditions under which a biological sample was taken from an Observation unit. A multitude of samples can be taken from a single observational unit. Each of these samples may also be subjected to one or more Assays.

To encourage domain researchers to FAIRify their data in the best possible way, a metadata registration tool should be flexible, and require little or no training. For the human readable high-level metadata registration, we have chosen for an intuitive web form. Next the tool prompts users to choose a minimal information model that best represents the type of sample (Figure 1). The chosen model specifies a set of mandatory and optional attributes that should be used to describe the samples taken. After selection of the most appropriate minimal information model(s) and relevant optional attributes, the tool will generate a metadata template workbook in an Excel format that will allow sample metadata registration in the form of attribute name-value pairs (Figure 2). Excel based template workbooks allow for offline, on-site metadata registration, support collaborative efforts, and information collection in high throughput.

**Figure 1.**
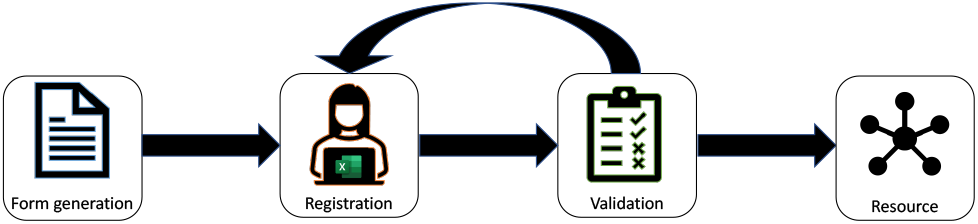
FAIR-Data station metadata registration workflow. The FAIR-Data station workflow consists of three main modules. Based on the minimal information check-list(s) selected by the user the “form generation module” creates a standardised metadata template excel workbook, the “validation module” checks the format of the metadata recorded in the workbook. The “resource module” exports the complete set of recorded metadata into an RDF data file, enabling (cross-project) meta data searches, and optionally into ENA compatible metadata submission files.

**Figure 2.**
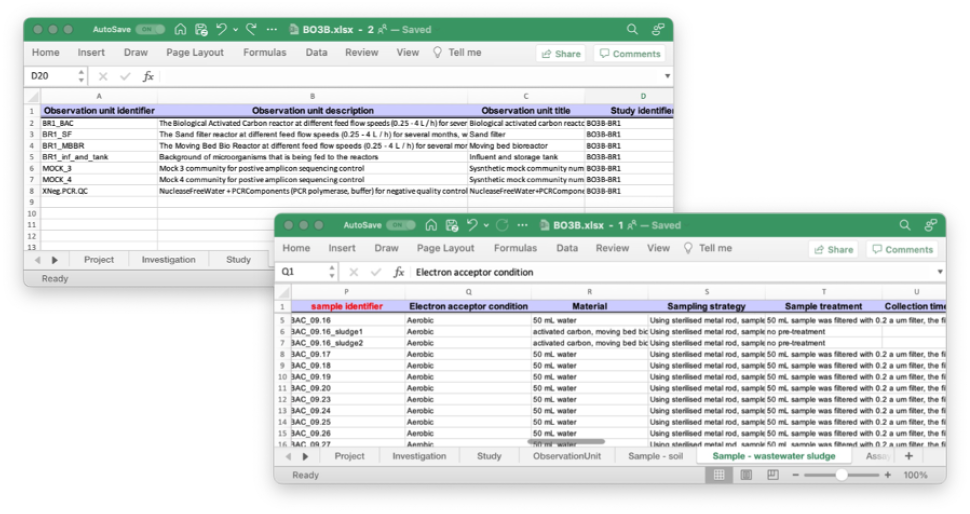
Observation Unit and Sample view of a project metadata workbook generated by the FAIR data station. Column headers represent the mandatory and optional attributes selected by the user. Each line represents the metadata values associated with a single sample. While the columns are in a default order, they can be rearranged to user’s preference and user-defined columns such as in this example “Electron acceptor condition” can be added. User-defined attribute-value pairs are not validated but user-defined column headers will be used as predicates in the RDF knowledge graph. Note that this a multi-sheet workbook in accordance with the ISA standard.

## Metadata selection and validation

To assist domain researchers in creating high-quality FAIR metadata the FAIR-DS comes with a library of 40 frequently used minimal information checklist: 23 are MIxS standards [10] not limited to sequence data and 17 are ENA minimal information checklists [11]. Each individual package contains a set of mandatory shared (core) attributes that should be included regardless of the chosen package. Checklist specific attributes are optionally selected by the user. This library is also a file in Excel format allowing researchers to easily add, update and extend existing standards, to develop new standards and change the pre-set status of optional and mandatory attributes. To be able to link different sample types to an observation unit and to be able to link multiple assay types to a sample, multiple checklists can be chosen in parallel which become available as individual sheets in the workbook.

To ease the hassles of creating high-quality metadata as much as possible, the FAIR-DS uses an Excel based template to register sample metadata. Excel files can be handled by many devices which opens the way for on-site metadata registration, for instance while taking a sample. During this process, recorded metadata values can be checked for having the correct format by simply uploading the Excel workbook to the FAIR-DS tool. There are several attributes with specified values. Boolean attributes for instance, should be true or false. All other values are invalid and using them compromises structural interoperability and therefore the machine-actionability of the metadata field. To warrant structural interoperability of recorded metadata, the uploaded Excel worksheet is validated by checking the format of testable values using regular expressions. Other checks include activation of unsolicited auto-complete and auto-corrections functions such as the transformation of a numeric value to a calendar date, and for mismatches between identifiers used at the different levels. In addition, we included support for reg-ular expressions such as “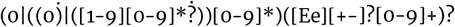 (glmLlmglng)” obtained from the ENA checklist [12] for sample volume or weight for DNA extraction.

## Querying metadata

Having experimental metadata at hand is key for efficient data analysis. After metadata validation an RDF file is automatically created based on the content provided. Multiple ontologies and terms are incorporated (FOAF, JERM, PPEO, PROV, Schema.org and MIxS) to generate an understandable resource of the experimental data [13,14, 9,15,16,10]. This file, in Turtle format, can be directly ingested in a triple store and creates the opportunity for researchers to directly query their metadata from different programming languages such as R, Python or Java and to incorporate the metadata in their analysis workflows. The impact of such a resource will become more significant if the FAIR Data Station is used in multiple research projects revolving around a common theme. Bringing together multiple project specific metadata (Turtle) files enables crosswalks between similar projects and questions such as “retrieve all samples of a specific type for which attribute X is “true”. Without a proper metadata management system such simple questions would be nearly impossible to ask.

## ENA submission of sequence files

One of the public resources to share and publish nucleotide data is the European Nucleotide Archive (ENA) as part of the ELIXIR infrastructure [17]. This resource is synchronised with public resources such as the National Center for Biotechnology Information (NCBI) ensuring that the research data is available from multiple sites. To convert research metadata into an ENA acceptable format, an ENA submission module was developed. This module accepts a validated RDF data file as input and converts study, observation unit, sample and assay metadata into XML files that can be directly uploaded to the ENA submission portal. Project PRJEB54921 is an example of such ENA submission. The metadata files used for this submission are available in the documentation.

## Implementation and Documentation

The FAIR Data Station (FAIR-DS) is a web-based Java application using Vaadin as a front end [18]. It is available as a JAR package and can be executed out-of-the-box without additional dependencies as a private or local instance. The FAIR-DS supports the FAIR-By-Design principles that aims to collect FAIR experimental metadata already from the first phase of a project.

Documentation is available as a Jupyter Book at http://fairbydesign.nl This includes technical information on how to set-up the FAIR Data Station, how to modify the already existing metadata schemas and how to extend this with new schemas. For users it is explained with telling examples in detail how to register and validate metadata, how to query the validated and converted data files and how to submit sequence related metadata to the ENA database.

## Conclusions

The FAIR Data Station is lightweight stand-alone application for metadata management and validation and was developed as an integral part for the UNLOCK infrastructure [https://m-unlock.nl] for exploring new horizons for research on microbial communities [19]. It has multiple features that enhance usability and interoperability: First, portability, the FAIR-DS is a stand-alone Java application including all dependencies. No additional installation steps are needed to use this program. Second is the usage of Excel Workbooks as a familiar environment for metadata registration. Out of the box Excel Workbooks provide multiple ways to present a clear overview of the metadata and enable cooperation and offline management. The use of an Excel Workbooks for sample registration separates the FAIR-DS from Dendro, CEDAR, *-DCC and COPO as these fairification tools are fully web-based [20, 21, 22, 23]. Lastly, the ability to automatically generate machine-actionable ENA metadata submission files will increase the FAIRness of sequence data submissions.

## Availability of source code and requirements

- Project name: FAIR Data Station
- Project home page: http://fairbydesign.nl
- Project git repository: https://gitlab.com/m-unlock/fairds
- Documentation: http://docs.m-unlock.nl
- Operating system(s): Platform independent
- Programming language: Java
- Other requirements: Java 11 or higher
- License: Apache License 2.0

